# Weakly-Supervised Prediction of Cell Migration Modes in Confocal Microscopy Images Using Bayesian Deep Learning

**DOI:** 10.1101/810473

**Authors:** Anindya Gupta, Veronica Larsson, Damian Matuszewski, Staffan Strömblad, Carolina Wählby

## Abstract

Cell migration is pivotal for their development, physiology and disease treatment. A single cell on a 2D surface can utilize continuous or discontinuous migration modes. To comprehend the cell migration, an adequate quantification for single cell-based analysis is crucial. An automatized approach could alleviate tedious manual analysis, facilitating large-scale drug screening. Supervised deep learning has shown promising outcomes in computerized microscopy image analysis. However, their implication is limited due to the scarcity of carefully annotated data and uncertain deterministic outputs. We compare three deep learning models to study the problem of learning discriminative morphological representations using weakly annotated data for predicting the cell migration modes. We also estimate Bayesian uncertainty to describe the confidence of the probabilistic predictions. Amongst three compared models, DenseNet yielded the best results with a sensitivity of 87.91% ± 13.22 at a false negative rate of 1.26% ± 4.18.

## 1. INTRODUCTION

An increased understanding of cancer cell migration regarding environmental factors and drug treatment may provide us with clues on how to reduce the risk of metastasis. With an automated and quantitative approach to analyze the cell migration process, we open up for large-scale systems microscopy experiments and drug screening. Here, we specifically focus on mesenchymal migration, where the cells adopt two distinct migration sub-modes: *continuous* and *discontinuous*, and can also switch between modes [1].

During live-cell imaging, cells are repeatedly exposed to light for capturing dynamic cellular responses over time. Light over-exposure can cause photo-toxicity, changing cells behavior and possibly cause apoptosis, and thereby reducing the number of cells for downstream experiments [1]. Therefore, we aim to determine individual cell migration modes from a minimal number of frames, capacitating gene expression analysis in single cells, disected from a cell culture.

The study of cell migration yields an overabundance of experimental data that requires demanding processing and human efforts. With an increasing amount of time-lapse sequences, the biologists are often unable to annotate (like in our case) the migration modes in each image-frame with high confidence, which makes the ground-truth labels noisy. Thus, it is desired for learning-based approaches to be able to work with weak supervision. Weak supervision provides a simple, model-agnostic way to integrate the domain-expertise into a learning model [2].

As in all applications of supervised deep learning, an insufficient understanding of model outputs may provide sub-optimal results [3, 4]. Estimating uncertainties (i.e., aleatoric and epistemic) can eventually increase the confidence for the predictions and lead to an improved decision [5]. Model uncertainty (epistemic) for an image can be obtained by keeping the dropout mechanism on at test time and performing multiple predictions using Monte Carlo sampling, which is similar to Bernoulli approximated variational inference [3, 5].

There are several existing methods, dealing with migration problem using cell tracking [6]. However, merely tracking will not reveal the migration mode as migration is heavily characterized by cell morphology. Recent studies have shown that the current cell morphology influences its future movement [7]. Therefore, we were inspired to explore whether convolutional neural networks (CNNs) can be exploited to predict migration modality based on the current cell morphology. In this study, we compare three popular CNN architectures for predicting the migration mode in a single cell frame. We employ Bayesian CNNs for probabilistic prediction of mesenchymal migration modes from weakly annotated data.

## 2. IMAGE DATA AND ANNOTATIONS

High resolution images (1024×1024×3 px) of H1299 human non-small lung carcinoma cells stained with EGFP-Paxillin (CMAC marker), RubyRed-LifeAct (F-actin marker) and a far-red membrane dye were acquired using Nikon A1R confocal microscope with 60× objective. The images were acquired for 8–10 hr at 5 min intervals with a pixel resolution of 0.21 *µ*m, resulting in 90-110 frames per time-lapse sequence. Altogether, the dataset consists of images with 1-3 cells, plated onto 96-well glass plates pre-coated with two different Fibronectin (FN) concentrations (10 *µ*g/ml, or *µ*g/ml). All the images were downsampled by a factor of two (512×512×3 px) to fit the networks (see Sect. 3).

An expert biologist annotated cells in each frame as either *discontinuous, continuous* mode or *unknown* for the frames difficult to visually interpret. The *unknown* class is discarded from both training and testing phases. After filtering out, we obtained 137 single cell sequences to train and evaluate our method. Amongst all, 78 and 54 cells are labeled as *continuous* and *discontinuous* modes throughout the time-lapse sequence whereas five cells switched between these modes.

We selected 102 sequences for training and 31 sequences for independent testing. The number of images is larger for one migration mode than others, presenting a class imbalance during training. We thus employ stratified sampling, ensuring that the relative class frequencies are approximately balanced in each fold. We also ensure that the images from the same cell are not present in both training and validation sets.

We augmented the training images by applying horizontal and vertical flipping, and multiple 90° rotations. The augmented images were further extended by including translation up to ± 2 pixels from the centroid position in both *x-* and *y-* directions, resulting in 72 variations per image.

## 3. PROPOSED METHOD

### Preprocessing

We extracted 227×227×3 px patches from the manually annotated centroid on each cell frame to include sufficient contextual information as an input for the network model. The input image intensities in all 3 channels are then normalized to zero mean and unit standard deviation.

### Weak supervision label assignment

Our annotations are noisy as manually characterizing the mesenchymal migration modes with high confidence is challenging due to their inherent switching properties. It is thus reasonable to predict the confidence of migration modes instead of directly classifying them. Motivated by [8], we introduce a weakly-supervised learning-based criterion. Unlike the former approach, we formulate the classification problem into regression by generating probabilistic labels from the discrete frame-wise labels.

We begin with an assumption that confidence in visual assessment increases over time when the cell behaves similarly throughout the sequence and decreases during switching behavior. As we are provided with discrete frame-wise labels of length *n’* for every cell sequence of length *n* in the training phase, the goal is to predict probabilistic confidence for the training data, as well as for unseen testing data. To achieve that we linearly interpolate the frame-wise labels in a continuous range of [0, 1] for assigning them to their corresponding *n’/n* frames. We linearly mapped labels of a cell corresponding to *continous* mode in the range 0 - 0.4 and *discontinous* mode in the range 0.6 - 1. The range between 0.41 - 0.59 characterizes the confidence regarding the conditions when a cell starts switching migration modalities.

### Bayesian Uncertainity

We measure uncertainty for each image sample *x* by independently dropping (with probability *P*_*drop*_) the weights in all layers while drawing Monte Carlo samples from a Bernoulli distribution. The predictive uncertainity is estimated by:

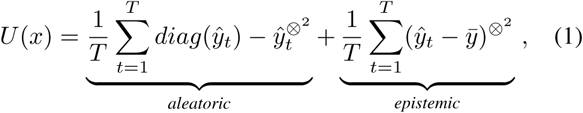

where 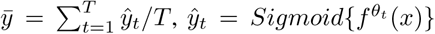 and *T* refers to as the sampling rate. We fixed *T*= 50, as it was found in our case to be sufficient for predictive mean estimation.

### Network models

We modified three existing networks, i.e., VGG16 [9], Resnet50 [10], DenseNet [11], to our need for comparison purposes. We transformed each network model to function as a variational dropout network [12], i.e., each weight of a model has a dropout rate, that allows efficient approximation of Bayesian inference. Each convolutional layer in all three models was employed with ℒ_2_ regularization to prevent overfitting, and is equivalent to putting a Gaussian prior on the network parameters, resulting in a maximum-aposteriori (MAP) solution [3].

We employed the batch normalization operation specifically in the VGG16 network model and replaced the fully-connected layers with global average polling operation to reduce the computational complexities. To obtain the regression output, we replaced softmax with sigmoid in the final activation layer of all three models. The dropout rate (*P*_*drop*_) for the initial layer in all three models was fixed to 0.1, which grows linearly at a rate of 0.02 for each subsequent layer. We found it to be a good compromise between getting a reasonable performance and uncertainty measures.

We trained all three models in a 5-fold grouped stratified cross-validation scheme for 30 epochs with an initial learning rate of 0.01 using Tensorflow backend in Keras, where the overall training took twenty hours on a Titan X GPU. The weights were initialized using Glorot normal distribution and the biases were set to zeros. The weights were updated in a batch of 32 samples using the ADAM optimizer. The networks were optimized by minimizing the logarithmic hyperbolic cosine error as loss function. All related codes are available at: 10.5281/zenodo.3490575

## 4. RESULTS

We present the mean square error (MSE) for all three models over 31 cell sequences from the independent test set in Fig. 1.a. Although all the three models performed reasonably well, we observed that the MSE monotonically reduces with increasing model capacity. DenseNet performed better than the other two models because of its inherent densely-connected skip connections, enabling it to capture fine morphologically discriminative representations.

**Fig. 1:**
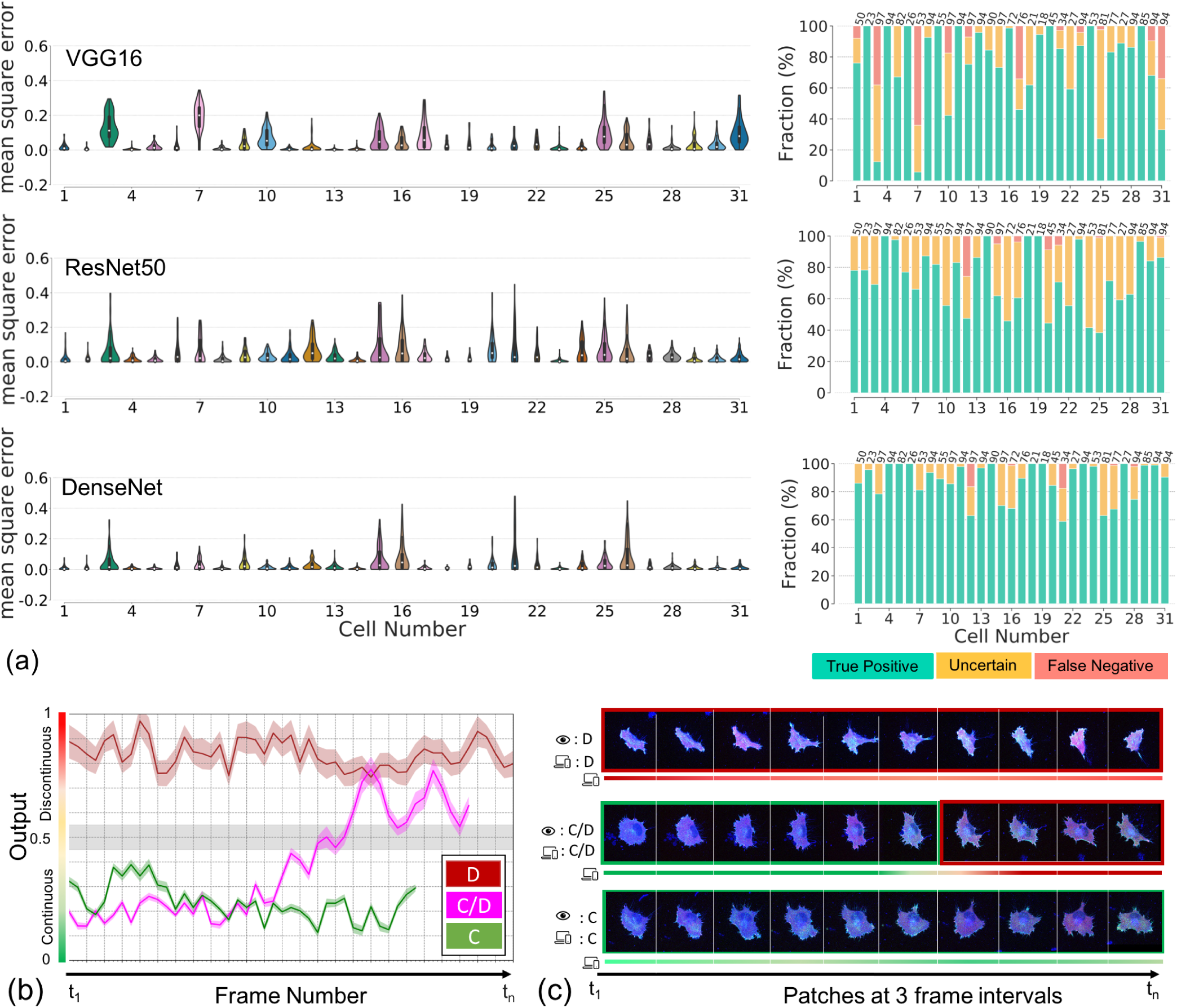
Comparison of models. (a) left: Violin plots of MSE per cell sequence, right: fraction (in %) of correctly classified cells per sequence. The number at the top corresponds to the number of frames in the sequence. (b) shows prediction results of the best CNN model with confidence intervals determined by epistemic uncertainty for three example cell sequnces. (c) shows visual results verification for these three cells at three frames interval are shown in (c). Here, C and D refer to only *continuous*, and *discontinuous* modes, and C/D mode to switching between them.

This observation is further validated by the results shown in the second column of Fig. 1.a. Here, we further quantitatively compare all three models by adopting prediction errors per frame as a metric to determine their confidence per image. In particular, we consider the result as true positive only when the predicted values are in the ranges of *continuous* and *discontinuous* as described in Sect. 3 and as Uncertain when the predictions range between 0.4-0.6. Each bar plot represents: *True positive* (TP) – correct prediction, *False negative* (FN) – wrong predictions, and *Uncertain* – less confident predictions; as the fraction of total number of frames in the cell sequence. Notice how the TP rate improves with increasing network capacity. Both Resnet50 and DenseNet models yielded reasonably similar performances; however, there were substantial differences in their confidence regarding predictions. Overall, DenseNet yielded the best results: a TP rate of 87.91% ± 13.22, and Uncertain rate of 10.83% ±11.28 at FN rate of 1.26% ±4.18.

According to both violin and bar plots, we can observe that misclassifications (FN) were more frequently seen when predictions are made using VGG, compared to that of Resnet50 and DenseNet. We assume that it was mainly because these models receive concatenated feature maps from the preceding layers as an input, allowing them to encode more diversified and richer representations. Also, VGG uses only the most complex features for predictions, whereas the other two models use representations of all complexity levels, thus providing smoother decision boundaries.

Detailed results for three test cells classified by DenseNet are shown in Fig. 1.b (displaying confidence intervals per cell and as well as per frame), and sample frames at three intervals are shown in Fig. 1.c. Both the human expert and CNN might, to a large extent, take into account the cell morphologies in the prediction process. However, CNNs are able to predict migration modes from a static frame because they can observe subtle morphological characteristics in cell migration modes that are somewhat difficult to notice for human-observers.

## 5. CONCLUSION

We present an automated approach for predicting the mesenchymal cell migration mode using weakly supervised data. Our results indicate that the CNNs can capture discriminative morphological representations directly from static images. To address the observer prediction variability, we modified and compared three popular CNNs by modeling Bayesian approximation into them for providing probabilities with model uncertainties that describe the prediction confidences. Our comparison study between the models with differential information supports the utility of the epistemic uncertainty. Computing epistemic uncertainty using Monte Carlo samples for variational inference is fast and can also be applied to already trained models.

In the future study, we will visualize the features of the cell images that were learned by the CNNs and contributed to their prediction (e.g., the protrusions and trailing edge) to reveal how and why CNNs can predict the migration modes. Given the spatiotemporal behavior of cells, we also intend to explore recurrent neural networks to encode morphological and temporal representations. For that, we are collecting more imaging data with annotations from multiple observers.

## 6. ACKNOWLEDGMENTS

This work is funded by the Swedish Foundation for Strategic Research grant SB16-0046. Sincere thanks to Ankit Gupta for his appreciative suggestions.

## Notes

https://github.com/anindgupta/isbi2020

